# The effects of aging and hearing impairment on listening in noise

**DOI:** 10.1101/2023.01.20.524873

**Authors:** Ádám Boncz, Orsolya Szalárdy, Péter Kristóf Velősy, Luca Béres, István Winkler, Brigitta Tóth

## Abstract

Listening in a noisy environment (e.g., speech in noise) relies on the fundamental ability to extract coherence from the variable sensory input. This allows the detection active sound sources and their segregation of them from the rest of the scene (figure-ground segregation). Peripheral and central causes of age-related decline of listening in noise were assessed by a tone-cloud-based figure detection task. In two conditions differing in the amount of noise, figure detection performance was equalized between young, normal-hearing, and hearing-impaired elderly listeners by adapting the stimulation separately to the abilities of each person. Based on behavioral measures and event-related brain potentials (ERP), in the absence of cognitive deficits, aging alone does not appear to significantly deteriorate the ability to detect sound sources in noise, although ERPs show delayed perceptual processes and some expected deterioration in attention and/or executive functions. However, even mild hearing impairment substantially reduces the ability to segregate individual sound sources within a complex auditory scene, and susceptibility to masking noise increases together with the severity of the hearing deficit.

**Significance Statement:** This work provides new information about the contributions of central and peripheral causes to the typical age-related decline of listening in a noisy environment. Behavioral and neurophysiological data collected in a well-controlled model of listening in noise suggest that aging alone does not significantly reduce the ability to detect sound sources in a complex auditory scene. However, even mild hearing impairment significantly reduces this ability. The stimulus paradigm used appears to be quite sensitive to hearing loss, making it potentially useful for the early detection of hearing problems.

## Introduction

In everyday situations, detecting auditory objects in noise (figure-ground segregation), such as understanding speech in a crowded restaurant, is essential for adaptive behavior. Difficulty in listening to speech under adverse listening conditions results in compromised communication and socialization, which are commonly reported phenomena in the aging population ((1), (2) (3), affecting ca. 40% of the population over 50, and almost 71% over 70 years of age (4, 5).

Problems in figure-ground segregation are attributed to peripheral, central auditory, and/or cognitive factors (5, 6). However, the relative contributions and interactions among these factors have not been studied in depth. Here we investigated together the effects of peripheral (hearing loss) and central auditory processes on the deterioration of figure-ground segregation in aging combining personalized psychoacoustics and electrophysiology.

### Aging and listening in noisy environments

Major changes in the peripheral auditory system likely contribute to age-related hearing loss. For instance, outer and inner hair cells within the basal end of the cochlea degrade at an older age, resulting in high-frequency hearing loss (5, 7). The pure-tone audiometric threshold is a proxy of changes in cochlear function and structure (5), and it is well established that due to elevated hearing thresholds elderly people have difficulty hearing soft sounds (8, 9). Age-related damage to the synapses connecting the cochlea to auditory nerve fibers (cochlear synaptopathy) results in higher thresholds with fibers having low spontaneous rates (10, 11), which is assumed to contribute to comprehension difficulties when speech is masked by background sounds (12). However, older adults with similar pure-tone thresholds can differ in their ability to understand degraded speech, even after the effects of age are controlled for (13).

Deficits in central auditory functions (decoding and comprehending the auditory message; (14, 15) may also contribute to the difficulties elderly people experience in speech-in-noise situations. Specifically, these central functions may partly compromise concurrent sound segregation (6, 16, 17) and lead to diminished auditory regularity representations and reduced inhibition of irrelevant information at higher levels of the auditory system (16, 18, 19).

Navigating noisy scenes also depends on selective attention (independently of the modality of stimulation), which is known to be impaired in aging (for reviews see (20, 21). Specifically, increased distraction by irrelevant sounds(1–3) suggest deficits in inhibiting irrelevant information, which is especially prominent in information masking(17).

### Behavioral and neural indices of figure-ground segregation of tone-cloud stimuli

Figure-ground segregation relies on grouping sound elements belonging to one sound source and segregating them from the rest of the competing sounds (22–25). Figure-ground segregation has been recently studied with the help of tone clouds, a series of short chords composed of several pure tones with random frequencies. The figure within the cloud consists of a set of tones progressing together in time, while the rest of the tones randomly vary from chord to chord (background; (24–27). When the frequency range of the figure and the background tone set spectrally overlap, the figure is only distinguishable by parsing the coherently behaving tones across frequency (concurrent grouping) and time (sequential grouping). With component tones of equal amplitude (which is typical in these studies), the ratio of the number of figure tones (figure coherence) and background tones (noise) determines the figure-detection signal-to-noise ratio. Figure detection performance within these tone clouds was found to predict performance in detecting speech in noise (12, 28), making this well-controlled stimulus attractive as a model for studying real-life figure detection.

Figure-related neural responses commence as early as after two temporally coherent chords (ca. 150 ms from the onset of the figure; (24, 27, 29). Figure detection accuracy scales with figure coherence and duration (the number of consecutive figure tone-sets presented). This suggests that both spectral and temporal integration processes are involved in figure detection (25). Event-related brain potential (ERP) signatures of figure-ground segregation are characterized by the early (200-300 ms from stimulus onset) frontocentral object-related negativity response (ORN); a later (450-600 ms), parietally centered component (P400) is elicited when listeners are instructed to detect the figure (25). The former indexes the outcome of the process separating the figure from the background (ORN is elicited when separating concurrent sound streams;(30, 31), while the latter likely reflects a process leading to the perceptual decision, such as matching to a memorized pattern (25, 30, 31).

While a mechanism based on tonotopic neural adaptation could explain the extraction of a spectrally coherent figure, it is more likely that this kind of figure detection is based on a more general process, such as the analysis of temporal coherence between neurons encoding various sound features(25, 26, 32). This is because spectrally constant and variable figures can be detected equally efficiently (24–26) and the segregation process is robust against interruptions (27).

Some results suggest that figure-ground segregation is primarily pre-attentive(26, 27). However, there is also evidence that figure-ground segregation can be modulated by attention (25) and cross-modal cognitive load (33).

### Research questions and hypotheses

The purpose of the current study was to test the causes of impaired listening in noise in aging. To separate the effects of age and age-related hearing loss, three groups of listeners (young adults; normal-hearing elderly, and hearing-impaired elderly) have been tested. The group of hearing-impaired elderly was selected based on an elevated pure-tone audiometric threshold, thus assuring deterioration of peripheral function (5). The effects of differences in peripheral gain and cognitive load (e.g., effects of the inter-individual variation in working memory capacity) between the groups were reduced by keeping task performance approximately equal across all listeners using individualized stimuli. Because performance in psychoacoustic measures inevitably combines the effects of peripheral and central processes, ERP measures (specifically ORN and P400) were collected to shed light on the underlying processes.

Participants were presented with stimuli concatenating 40 chords of 50 ms duration (Figure 1A). Half of the stimuli included a figure (a set of tones rising together in time embedded in a cloud of randomly selected tones; Figure trials), while the other half consisted only of chords made up of randomly selected frequencies (No-Figure trials; stimuli were adapted from (24–26)). Listeners performed the figure detection task under low noise (LN) and high noise (HN) conditions, which differed only in the number of concurrent randomly varying (background) tones. Stimulus individualization was achieved through two adaptive threshold detection procedures conducted before the main figure detection task. First, low-noise stimuli were adjusted for each participant by manipulating the number of tones belonging to the Figure (figure coherence) so that participants performed figure detection at 85% accuracy. Second, high-noise stimuli were adjusted for each participant by increasing the number of background tones in the LN stimuli until the participant performed at 65% accuracy.

**Figure 1.**
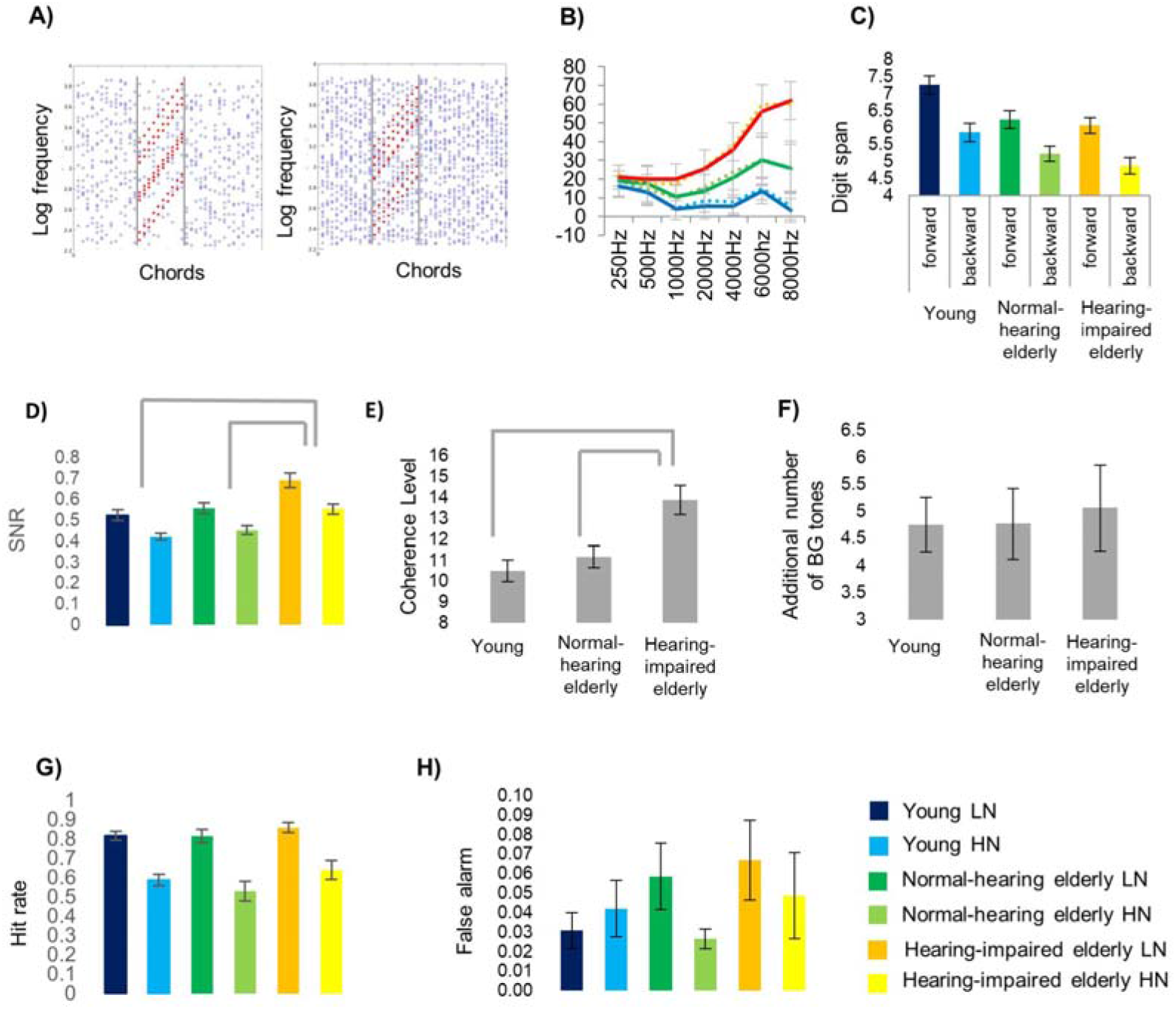
(A) Figure Examples of stimuli for the low (LN) and high noise (HN) conditions, respectively. (B) Mean pure tone audiometry thresholds for the young, adult (blue line; N=20), normal-hearing, elderly (green line; N=13), and hearing-impaired older listeners across elderly groups (red line; N=16) in the 250 - 8000 Hz range. Error bars depict SEM on all graphs. (C) Forward and backward digit span performance (corresponding to working memory capacity and control) for the three groups. (D) Mean SNR values were derived from the threshold detection tasks ((across LN and HN) from the stimulus individualization procedure. (E) Mean figure coherence level derived from the first threshold detection task (and employed later in the FG segregation task) for the three groups. Mean Figure coherence level of the LN stimuli for the three groups.; significant group differences (p<0.01) are marked by gray lines above the bar charts. (F) Mean increase in the background tone number between the tones from LN to HN conditions for the three groups. (G-H) Behavioral performance (Hit rate and false alarm rate, respectively) in, separately for LN and HN; color labels are at the FG segregation task lower left corner of the figure.

If integrating sound elements into a single object was more difficult for the elderly than the young listeners, then they will require more figure tones than young listeners to reach similar figure detection performance, as higher coherence helps figure integration (24–26). Suppose the elderly adults were more susceptible to masking. In that case, their performance will be affected by fewer additional background tones than young adults when individual stimuli are created for the HN condition.

As for separating peripheral and central processes, peripheral effects are expected to be more pronounced in the elderly group with hearing loss than in the normal-hearing elderly: higher coherence and/or fewer background tones are needed for reaching the same performance. In contrast, both groups will be equally affected by the deterioration of central processes. One general effect often seen in aging is slowing information processing (20, 21, 34). One should expect longer ORN or P400 latency in both elderly groups compared to young adults. Finally, less efficient selective attention may lead to lower P400 amplitudes.

## Results

### Behavioral results

Participants were divided into three groups based on their age and pure-tone hearing threshold (Figure 1B), the latter measured by audiometry (young adults, normal-hearing elderly, and hearing-impaired elderly). None of the listeners reported issues with their hearing (“the hearing handicap inventory for the elderly” adopted from (35). After measuring the participant’s digit span and introducing the stimuli in general, the LN and HN conditions were set up individually for each participant. First, the number of parallel figure tones allowing the participant to detect the figure with 85% accuracy was established by a procedure of increasing the number of figure tones while simultaneously decreasing the number of background tones, thus keeping the total number of tones constant (*N* = 20; LN condition). Then, extra background tones were added, until the participant’s performance declined to 65% (HN condition). The main experiment consisted of a series of trials mixing LN and HN, Figure, and No-Figure stimuli in equal proportion (25%, each). Figure detection responses and EEG were recorded. Figure 1 summarizes behavioral results from the pure tone audiometry, digit span, and FG segregation task.

#### Group differences in the number of figures and background tones in the LN and HN stimuli

The threshold detection procedure yielded distinct LN and HN conditions for each listener. The signal-to-noise ratio (SNR) was calculated from the ratio of the number of figure tones and background tones, separately for each group and the LN and HN conditions (Figure 1D). SNR values were log-transformed before analyses due to their heavily skewed distribution. As was set up by the procedure, there was a main effect of NOISE (LN vs HN conditions) on SNR (*F*(1, 46) = 75.57, *p* < 0.001, *n_p_^2^* = 0.622), with larger SNR in the LN (before log transformation: *M* = 2.29, *SD* = 3.60) than in the HN (*M* = 0.99, *SD* = 0.53) condition.

There was also a main effect of GROUP: *F*(2, 46) = 10.93, *p* < 0.001, *η_p_^2^* = 0.697. Post-hoc pairwise comparisons revealed that SNR was larger in the hearing-impaired elderly group (*M* = 2.85, *SD* = 4.36) compared to both the normal-hearing elderly (*M* = 1.12, *SD* = 0.54; Tukey’s HSD *q*(2, 95) = 5.35, *p* < 0.001) and the young adult group (*M* = 1.00, *SD* = 0.54; *q*(2, 95) = 7.30, *p* < 0.001). The interaction between GROUP and NOISE showed a tendency towards significance (*F* (2, 46) = 2.86, *p* = 0.067, *n_p_^2^* = 0.111). Separate pairwise comparisons between the two NOISE levels for each group showed that the effect size of NOISE was smaller in the hearing-impaired elderly than in the other two groups (all *p*s < 0.001; effect size for young adults: Cohen’s *d* = 1.78; for normal-hearing elderly: *d* = 1.98; for hearing-impaired elderly: *d* = 1.12).

To test whether the SNR effects were due to the number of coherent tones in the figure (coherence level; Figure 1G) or to the noise increase for the HN condition, separate ANOVAs were calculated for the coherence level (in LN) and the additional number of background tones (HN; Figure 1H). For coherence level, a one-way ANOVA with factor GROUP found a main effect (*F*(2, 46) = 9.62, *p* < 0.001, *η_p_^2^* = 0.295), with larger values in hearing-impaired elderly (*M* = 13.88, *SD* = 2.80) than in the normal-hearing elderly (*M* = 11.15, *SD* = 1.86; *q*(2, 95) = 4.57, *p* = 0.006) or young adults (*M* = 10.5, *SD* = 2.28; *q*(2, 95) = 5.67, *p* < 0.001). For the number of additional background tones, a one-way ANOVA with factor GROUP yielded no significant effect (*F* (2, 46) = 0.07, *p* = 0.93, *η_p_^2^* = 0.003).

#### Figure detection results

The effect of GROUP on task performance (*d*’, hit rate, false alarm rate, and RT; Figure 1E and F) was tested with one-way ANOVAs, separately for LN and HN. As was set up by the individualization procedures, there was no significant main effect of GROUP for any of the performance measures in the main figure detection segregation task in either noise condition (all *F*s(2, 46) < 1.67, *p*s > 0.2, *η_p_^2^*’s < 0.07).

#### Relationship between peripheral loss and figure-ground segregation

The relationship between peripheral hearing loss (average hearing threshold across frequencies and ears, as measured by pure-tone audiometry) and behavioral performance (log-transformed SNRs from the individualization procedure, as well as their difference; *d*’ and RT from the main task, separately for LN and HN) was tested with Pearson’s correlations. There were significant correlations between peripheral loss and SNR both in LN (*r*(47) = 0.556, *p* < 0.001; Bonferroni correction applied for all correlation tests) and HN (*r*(47) = 0.561, *p* < 0.001), as well as their difference (*r*(47) = −0.385, *p* = 0.044). The latter showed that larger peripheral loss (worse hearing) resulted in smaller noise differences between LN and HN. Confirming the success of stimulus individualization, no significant correlation was observed between peripheral loss and figure detection performance (*d’* and RT in either LN or HN; all *rs*(47) < 0.11, all *p*s > 0.5).

#### Relationship between working digit span and figure-ground segregation

The relationship between the digit span measures (working memory capacity and control, as measured by forward and backward digit span, respectively; Figure 1C) and behavioral performance (log-transformed SNRs of the individualization procedure and their difference; *d*’ and RT from the main task, separately for LN and HN) was not significant (all *r*s(47) < 0.265, *p*s > 0.5).

### ERP results

ERPs were separately collected for Figure and No-Figure trials, the LN and HN conditions, and the young adult, normal-hearing elderly, and hearing-impaired elderly group. Only ERPs to Figure and No-Figure events with a correct response (hit for Figure, correct rejection for No-Figure) were analyzed. Two ERP components were identified based on a visual inspection of the group’s average responses. Figure detection elicited the ORN response (Figure 2A and B) over fronto-central leads, followed by a parietally maximal P400 response (Figure 3A and B). Two contrasts were tested by mixed-mode ANOVAs on the amplitudes and latencies of the ORN and P400 responses with factors of FIGURE (Figure vs No-figure), NOISE (LN vs. HN), LATERALITY (left vs midline vs right), and GROUP: one for exploring age effects by comparing the young adult and the normal-hearing elderly group (AGE factor), and one to test the effect of age-related hearing loss by comparing the normal-hearing and the hearing-impaired elderly group (HEARING IMPAIRMENT factor). Effects not including GROUP (AGE or HEARING IMPAIRMENT) or NOISE are reported in the Supplementary Materials (SM).

**Figure 2.**
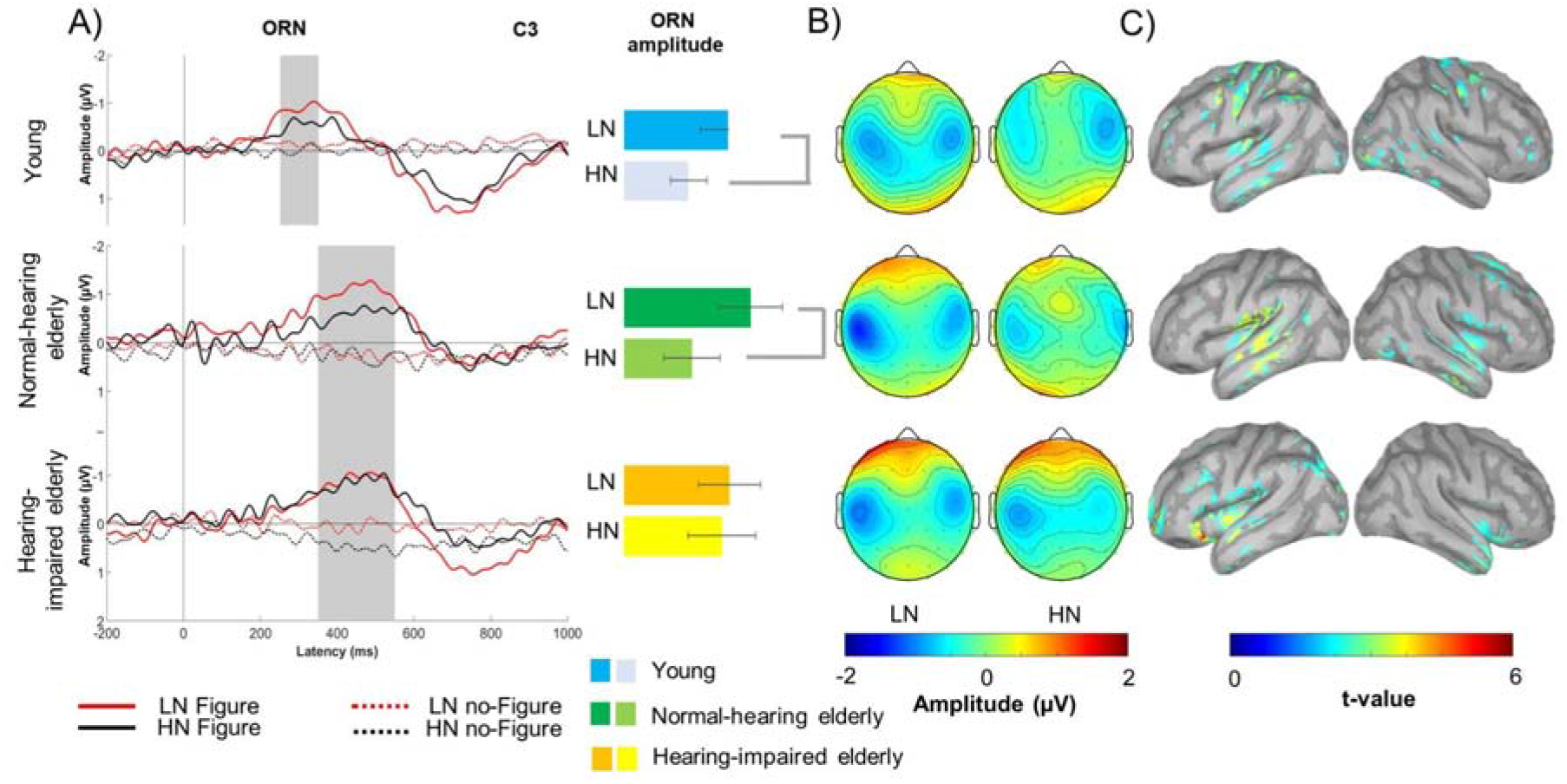
**A)** Group-averaged (young adult: N=20; normal-hearing elderly: N=13; hearing-impaired older elderly: N=16) central (C3; maximal ORN amplitude) ERP responses to Figure (solid line) and noNo-Figure (dashed line) related central (C3 lead) ORN elicited stimuli obtained in the LN (red) and the HN condition (black) respectively for young normal-hearing and hearing-impaired older listeners.). Zero latency is at the onset of the Figure event. Gray vertical bands show the measurement window for ORN while the yellow dashed line indicates the latency. The bar charts on the right side of ORN in young adults. On the panel shown on the right, the effect of NOISE is shown on the barplot for mean ORN amplitude respectively for amplitudes (with SEM) separately for the LN and HN conditions and groups. Significant NOISE effects (p<0.05) are marked by gray lines beside the bar charts. **B)** Scalp distribution of the ORN responses to Figure elicited ORN response respectively stimuli for the LN and HN conditions and groups with color scale below. C). Source localization results of) Brain areas sensitive to the NOISE effect (HN vs. LN condition) within the ORN time window. **C)** Significant NOISE effect on source activity (current source density based on dSPM) found in young normal-hearing and hearing-impaired older adult groups separately. (color scale below).

**Figure 3.**
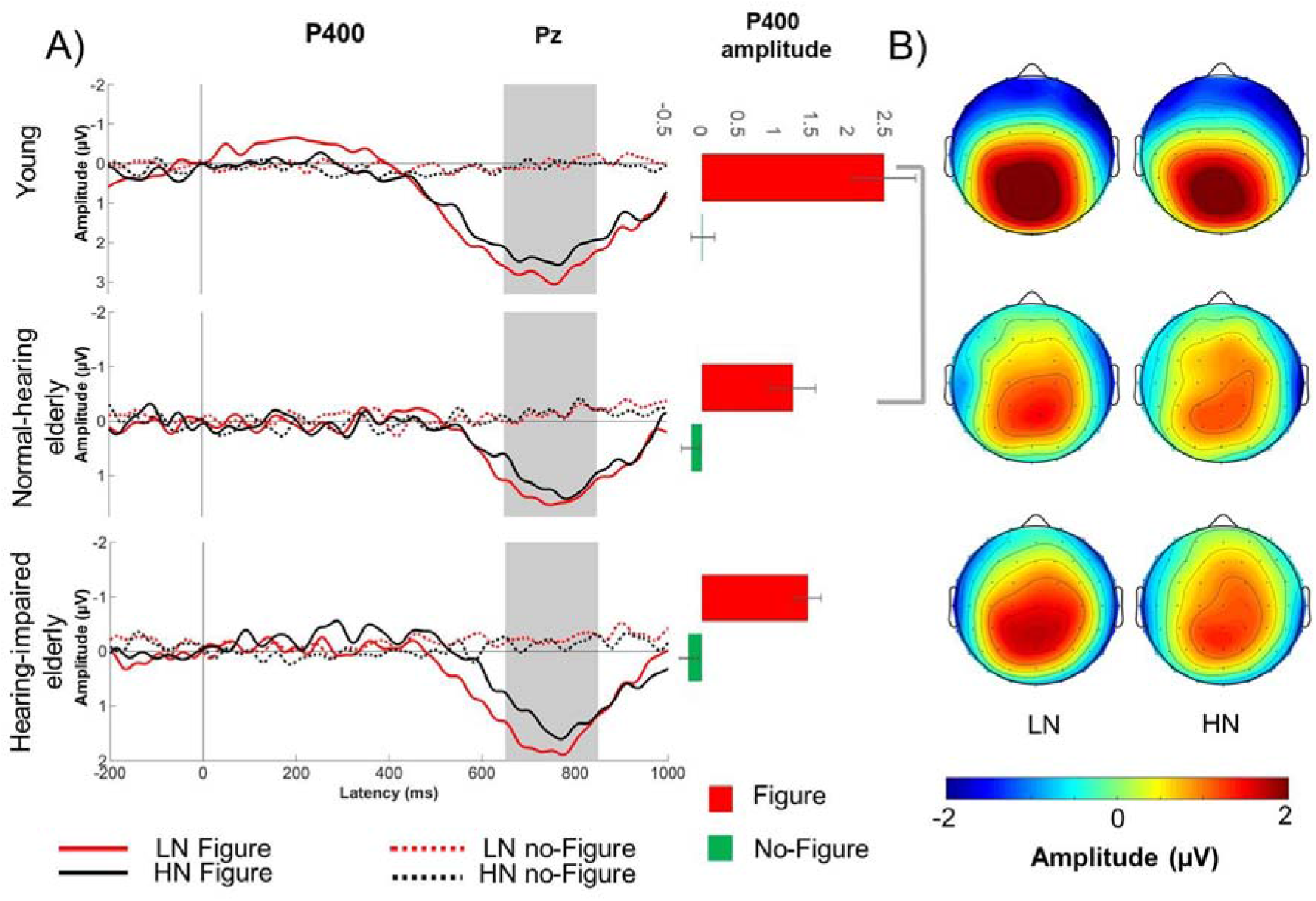
A) Group-averaged (young adult: N=20; normal-hearing elderly: N=13; hearing-impaired older elderly: N=16) parietal (Pz; maximal P400 amplitude) ERP responses to Figure (solid line) and No-Figure (dashed line) related parietal (Pz lead) p400 elicited stimuli obtained in the LN (red) and the HN condition (black) respectively for young normal-hearing and hearing-impaired older listeners.). Zero latency is at the onset of the Figure event. Gray vertical bands show the measurement window for P400. On the right, the effect of FIGURE is shown The bar charts on the barplot for the right side of the panel show the mean P400 amplitude respectively for Figure and no-Figure trial and groups. Amplitudes (with SEM) separately for the LN and HN conditions. Significant group effects (p<0.05) are marked by gray lines beside the bar charts. B) Scalp distribution of Figure elicited the P400 response respectively responses to Figure stimuli for the LN and HN conditions and groups with color scale below.

#### Object Related Negativity (ORN)

##### AGE effect on the ORN amplitude

No main effect of AGE was found on the ORN amplitude. There was a tendency for the NOISE effect (*F* (1, 31) = 4.069, *p* = .0524; *η_p_^2^* = 0.116) with larger (more negative) ORN amplitudes in the LN compared to the HN condition. The interaction between NOISE and FIGURE also yielded a tendency (*F* (1, 31) = 4.1550, *p* = .05012; *η_p_^2^* = 0.118) with Figure trials eliciting larger ORN for LN than HN (post hoc comparison: *p* = 0.032) but not No-Figure trials.

##### HEARING IMPAIRMENT effect on the ORN amplitude

The three-way interaction of FIGURE x NOISE x HEARING IMPAIRMENT was significant (*F* (1,27) = 4.942, *p* = 0.035; *η_p_^2^* = 0.15). This effect was further analyzed with repeated measures ANOVAs including only Figure trials measured at the C3 electrode (due to the left central ORN distribution; see SM), separately for the normal-hearing and hearing-impaired elderly groups. In the normal-hearing elderly group, Figure stimuli elicited significantly larger ORN in the HN relative to LN condition (*F*(1, 12) = 5.344, *p* = 0.039), whereas no significant NOISE effect was found in the hearing-impaired elderly group (*F*(1, 15) = .078, *p* = 0.783).

##### AGE and HEARING IMPAIRMENT effects on the ORN peak latency

Figure events elicited ORN between 250 and 350 ms latency from Figure onset in the young adult group while between 350 and 550 ms in the normal-hearing and hearing-impaired elderly groups. There was a significant main effect of AGE *F*(1,31) = 4.2662, *p* < 0.05, *η_p_^2^*= 0.12), with the ORN peak latency delayed in normal-hearing elderly (*M* = 421 ms) compared to young adults (*M* = 280 ms). NOISE and HEARING IMPAIRMENT did not significantly affect the ORN latency.

The brain regions activated during the ORN period were identified by source localization performed on the responses elicited by Figure trials. The sensitivity to signal-to-noise ratio was tested by comparing the source signals between the LN and HN conditions, separately for the young adult, normal-hearing, and hearing-impaired elderly groups by permutation-based t-tests. Significant NOISE effects were found predominantly in higher-level auditory and associational areas such as the left temporal cortices, the planum temporale (PT), and the intraparietal sulcus (IPS) Figure 2C). In young listeners, precentral cortical regions were also significantly sensitive to the signal-to-noise ratio.

#### P400

##### AGE and HEARING IMPAIRMENT effects on the P400 amplitude

The P400 amplitude was significantly lower in the normal-hearing elderly group compared to the young adults (*F*(1,31) = 5.00, *p* < 0.05, *η_p_^2^* = 0.14). A significant interaction was found between FIGURE and AGE (*F*(1,31) = 6.50, *p* < 0.05, *η_p_^2^* = 0.17). Post hoc comparisons revealed that the P400 response was lower in the normal-hearing elderly relative to young adults for the Figure responses (*p* = 0.008) but not for the No-Figure responses (*p* > 0.05). No significant effect including NOISE or HEARING IMPAIRMENT was found for P400 amplitude.

##### AGE and HEARING IMPAIRMENT effects on the P400 peak latency

Neither NOISE nor AGE or HEARING IMPAIRMENT significantly affected the P400 latency.

## Discussion

We tested the contributions of peripheral and central auditory processes to age-related decline of hearing in noisy environments using a tone-cloud-based figure detection task. We found that while aging slows the processing of the concurrent cues of auditory objects (long ORN latencies in the elderly groups) and may affect processes involved in deciding the task-relevance of the stimuli (lower P400 amplitude in the normal-hearing elderly than the young adult group), overall, it does not significantly reduce the ability of auditory object detection in noise (no significant differences in the signal-to-noise ratios between the young adults and the normal-hearing elderly group). However, when aging is accompanied by higher levels of hearing loss, grouping concurrent sound elements suffers and, perhaps not independently, the tolerance to noise decreases, as higher coherence was needed by the hearing-impaired than the normal-hearing elderly group for the same figure detection performance, hearing thresholds negatively correlated with the number of background tones reducing detection performance from the LN to the HN level, and the ORN amplitudes did not significantly differ between the HN and LN condition in the hearing-impaired elderly group (while they differed in the other two groups). The inference about the effects of peripheral hearing loss is strongly supported by the efficacy of the stimulus individualization procedure: there are no significant group differences and no significant correlation between hearing thresholds and performance measures of the figure detection task, and no correlation between the current working memory indices and any of the behavioral measures. Further, the current results are fully compatible with those of prior studies showing that spectrotemporal coherence supports auditory stream segregation (better figure detection performance with higher coherence(31,36–39).

We now discuss in more detail the general age-related changes in auditory scene analysis followed by the effects of age-related hearing loss.

### Age-related changes in auditory scene analysis

We found no behavioral evidence for age-related decline either in the ability to integrate sound elements (coherence level) or in the sensitivity to the noise (number of added background tones). Evidence about aging-related changes in auditory scene analysis is contradictory. Results are suggesting that the ability to exploit sequential stimulus predictability for auditory stream segregation degrades with age (40). A recent study, however, suggested that elderly listeners can utilize predictability, albeit with a high degree of inter-individual variation(37). Further, de Kerangal and colleagues (41)also found that the ability to track sound sources based on acoustic regularities is largely preserved in old age. The current results strengthen this view, as figures within the tone clouds are detected by their temporally coherent behavior. Further, age did not significantly affect performance in an information masking paradigm (42), a result fully compatible with the current finding of no significant behavioral effect of age, as tone clouds impose both energetic and informational masking on detecting figures.

Although figure detection performance was found preserved in the normal-hearing elderly group, the underlying neural activity significantly differed compared to the young adults both at the early (ORN) and later (P400) stage of processing Figure trials.

#### ORN results

In line with our hypothesis, the early perceptual stage of central auditory processing was significantly slowed in the elderly compared to young adults (the peak latency of ORN was delayed by ca. 150 ms). Although using the mistuned partial paradigm, Alain and colleagues ((6, 43) did not report a significant delay of ORN in the elderly compared to young or middle-aged adults, a tendency of longer ORN peak latencies with age can be observed on the responses (see (43) Figure 4). ORN is assumed to reflect the outcome of cue evaluation, the likelihood of the presence of two or more concurrent auditory objects (44). Compared to the mistuned partial paradigm, in which only concurrent cues are present (i.e., there is no relationship between successive chords), the tone-cloud-based figure detection paradigm also includes a sequential element: the figure only emerges if the relationship between elements of successive chords is discovered. It is thus possible that the delay is due to slower processing of the temporal aspect of the segregation cues. Alternatively, the delay may be related to the higher complexity of concurrent cues in the current compared to the missing fundamental design, because the latter can rely on harmonicity, whereas the figure in the current paradigm links together tones with harmonically unrelated frequencies. Slower sensory information processing has often been found in the elderly compared to young adults (e.g., Alain and McDonald found an age-related delay of the latency of the P2 component).

A possible specific explanation of the observed aging-related delay of the ORN peak is that the auditory system at older age needs to accumulate more sensory evidence for the perceptual buildup of the object representation. The input from the periphery may be noisier at an older age (for review see (5) therefore more time is needed to evaluate the relations between the current and previous chords or separate and integrate the spectral elements into an object than at a young age. This assumption is compatible with results showing that elderly listeners perform at a higher level in detecting mistuned partials with chords of 200 ms duration, compared to 40 ms duration and the resulting ORN responses are very similar to those obtained for young adults (6). Concordantly some studies using speech in-noise tasks suggested that older listeners required more time than younger listeners to segregate sound sources from either energetic or informational maskers (17, 45) The study from Ben-David et al demonstrated similar results for a speech babble masker but not for a noise masker (17).

#### P400 results

The P400 amplitude was significantly lower in normal-hearing elderly compared to young adults. Considering the commonalities between the neural generators and sensitivity to stimulus and task variables between the P400 and the P3 components (16, 20, 46, 47) P400 likely reflects attentional task-related processes (48). The P3 amplitude was found to be lower in healthy aging (49, 50). This is interpreted as normal cognitive decline with aging (49, 51, 52). Therefore, the current finding of reduced P400 amplitude likely reflects general cognitive age-related changes in attention or executive functions.

Distraction by irrelevant sounds (1–3) may be an important cause of the difficulties encountered by many elderly people in speech-in-noise situations. Deterioration of these central functions may partly compromise concurrent sound segregation (16, 17, 43), lead to diminished auditory regularity representations at the higher stations of the auditory system (18, 53), as well as deficient inhibition of irrelevant information processing (19).

### Consequences of age-related hearing loss on stream segregation ability

The hypothesis suggesting that integrating sound elements into an object was more difficult for the elderly with moderate hearing loss than for normal-hearing elderly was confirmed: hearing-impaired elderly needed more figure tones and higher SNR than normal-hearing elderly listeners to reach similar figure-detection performance. Specifically, while for normal-hearing elderly listeners ca. 55% of the tones in the chord forming the figure was sufficient for an 85% figure-detection ratio (LN condition) hearing-impaired elderly needed ca. 70% of the tones to belong to the figure for the same performance. Further, hearing impairment may increase susceptibility to masking by the background tones, as the number of background tones reducing performance from 85 to 65% (HN condition) negatively correlated with the hearing threshold.

The ORN responses may provide further insight into the problems of figure-ground segregation caused by hearing impairment. Whereas young adults and normal-hearing elderly elicited larger ORNs in the low than the high noise condition, the ORN amplitude did not differ between the two conditions for the hearing-impaired elderly. This suggests that the sensory-perceptual processes involved in detecting figures in noise were not made more effective by surrounding the figure with less noise for the hearing-impaired elderly. The lower amount of information arriving from the periphery limits their ability to find coherence or integrate concurrent sound elements. Consequently, there is no capacity to reduce the effect of additional noise resulting in a steep performance decline when in more noisy situations.

In contrast to early (sensory-perceptual) processing, no difference was found between normal-hearing and hearing-impaired elderly listeners, as was shown by the similar-amplitude P400 responses in the two groups. Thus, hearing deficits without general cognitive effects (as was promoted by the group selection criteria and the lack of working memory differences found between the group) only affect early sensory-perceptual processes. Further, as the level of hearing impairment of the current hearing-impaired group is modest (as none of the participants reported serious difficulties in the hearing handicap inventory for the elderly; (35)) the current results suggest that the tone-cloud-based figure detection paradigm could be used to detect hearing loss before it becomes severe.

## Conclusions

Results obtained in a well-controlled model of the speech-in-noise situation provided evidence that 1) age alone does not diminish the ability to detect auditory objects within a noisy scene and 2) age-related difficulties in listening under adverse conditions are in large part due to hearing impairment, which makes figure-ground segregation especially difficult for elderly people.

The tone-cloud-based figure detection task allows one to detect the decline of hearing before it becomes clinically significant.

## Materials and Methods

### Participants

Twenty young (14 females; mean age: 21.2 ± 2,4) thirteen old (9 females; mean age: 67.3 ± 4,3) with intact hearing and sixteen old (11 females; mean age: 68.7 ± 4,0) with mild hearing impairment participated in the study. Participants were financially compensated for their participation (ca. 4 euros/hour). None of the participants reported any neurological diseases. Participants gave informed consent before the experiment. Study protocols adhered to the Declaration of Helsinki and were approved by the local review board of the Institute of Cognitive Neuroscience and Psychology at the Research Centre for Natural Sciences.

#### Audiometry

All participants underwent standard pure tone audiometric hearing tests in which hearing thresholds (HT) were screened at frequencies between 250 Hz and 8000 Hz (250, 500, 1000, 2000, 4000, 6000, 8000 Hz), separately for both ears. Based on the hearing test results, elderly participants were divided into two groups: elderly participants with any HT above 45 dB were categorized as having mild hearing loss (5), while those with no HT above 45 dB formed the normal hearing elderly group. The threshold difference between the two ears was less than 20 dB in all participants. Average HT across tested frequencies was entered into a mixed-model ANOVA with between-subject factor GROUP (young, normal-hearing, and hearing-impaired elderly) and within-subject factor EAR (left or right). As expected, there was a significant main effect of GROUP (*F*(2, 46) = 101.18, *p* < 0.001, *η_p_^2^* = 0.788), but no main effect of the EAR (*F*(1, 46) = 1.39) or interaction (*F*(2, 46) = 0.83). Pairwise comparisons revealed that all three groups differed significantly in terms of average HT. Young adults had a lower average HT (*M* = 9.69 dB, *SD* = 2.97) than normal-hearing (*M* = 19.75 dB, *SD* = 6.80; *t*(31) = 4.87, *p* < 0.001, *d* = 0.99) and hearing-impaired elderly (*M* = 34.31 dB, *SD* = 5.81; *t*(34) = 16.50, *p* < 0.001, *d* = 2.55). Normal-hearing elderly also had a significantly lower HT than hearing-impaired elderly (*t*(27) = 6.22, *p* < 0.001, *d* = 0.93).

#### Subjective hearing impairment questionnaire

Subjective hearing impairment (i.e., hearing difficulties in everyday life including their emotional and social aspects) was assessed by a subset of the “Hearing Handicap Inventory for the Elderly” questionnaire (HHIE, adopted from (35); see the administered subset in S1 Table). Scores in the short HHIE test did not differ across groups (F(2, 42)=0.754, p=0.47).

#### Working memory assessment

Working memory (WM) capacity was tested by the forward and backward digit span tasks (54); see Figure 1) and compared across groups by one-way ANOVAs with the between-subject factor GROUP. Both the WM capacity (forward digit span) and WM control function (backward digit span) were significantly higher in young adults than in either elderly group (pairwise all cases p<0.005; forward digit span: F(2,46) = 6.694, p < 0.05, *η_p_^2^* =0.225; backward digit span: F(2,46) = 3.99, p < 0.05, *η_p_^2^* =0.148). No significant difference was found for either WM function between the two elderly groups of participants.

### Stimuli

Sounds were generated with Matlab (R2017a, Mathworks; Natick, MA, USA) at a sampling rate of 44.1 kHz and 16-bit resolution. Stimuli were adapted from previous studies (25, 26)samples depicted in Figure 1A). Each stimulus consisted of 40 random chords (sets of concurrent pure tones) of 50 ms duration with 10 ms raised-cosine ramps and zero inter-chord interval (total sound duration: 2000 ms). Each chord in the low-noise condition was composed of 20 pure tones with the tone frequencies drawn with equal probability from a set of 129 frequency values equally spaced with one semitone step in the 179-7246 Hz interval (the “tone cloud” stimulus). (See the number of concurrent tones in the high-noise condition in the “Threshold detection setting up the low and the high noise conditions” subsection.) A correction for equal loudness of tones with different frequencies was applied to the stimuli, based on the equal-loudness contours specified in the ISO 226:2003 standard.

In half of the stimuli (Figure stimuli), a subset of the tones increased in frequency by one semitone over ten consecutive chords. Thus, these tones were temporally coherent with each other, forming a figure within the tone cloud stimulus stimuli. Listeners can segregate temporally coherent tones from the remaining ones; perceiving this “figure” against the background of concurrent tones become more likely by increasing the number of temporally coherent tones (termed “coherence level”;(25, 26). In the current study, the figure appeared one for 500 ms randomly within the 300-1700 ms interval from stimulus onset (between the 7th and the 33rd position of the sequence of 40 chords). In the other half of the stimuli (No-figure stimuli) all tones of the chords were selected randomly.

### Procedure

Participants were tested in an acoustically attenuated and electrically shielded room at the Research Centre for Natural Sciences. Stimulus presentation was controlled with Psychtoolbox 3.0.16 (55). Sounds were delivered to the listeners via HD600 headphones (Sennheiser electronic GmbH & Co. KG) at a comfortable listening level of 60–70 dB SPL (self-adjusted by each listener). A 20” computer screen was placed directly in front of participants at 80 cm for displaying visual information.

#### Familiarization

At the beginning of the experiment, participants were familiarized with the stimuli by asking them to listen to Figure and No-figure stimuli until they were confident in perceiving the difference (~5 mins). To facilitate detection during familiarization, Figure stimuli were generated with a coherence level of 18 (out of the total 20 tones making up the chords). In each trial, the participant initiated the presentation of either a Figure or a No-figure stimulus by pressing a key on the left or the right side of a keyboard. The keys used in this phase were then consistently used for the Figure or the No-figure responses throughout the experiment; they were counterbalanced across participants. During familiarization, a fixation cross was shown at the center of the computer screen, and participants were instructed to focus on it. The familiarization phase ended when participants indicated their confidence in being able to distinguish between Figure and No-figure stimuli.

#### Training

Next, participants were trained in the Figure detection task (~10 mins). Each trial started with the presentation of a stimulus (2000 ms) followed by the response interval (maximum of 2000 ms), visual feedback (400 ms), and an inter-trial interval (ITI) of 500-800 ms (randomized). Participants were instructed to press one of the previously learned responses during the response interval to indicate whether they detected the presence of a Figure or not (2-alternative forced-choice detection task). The instruction emphasized the importance of confidence in the response over speed. During the ITI and the stimulus, a fixation cross was shown centrally on the screen, which then switched to a question mark for the response interval. Feedback was provided in the form of a short text (“Right” or “Wrong”) displayed centrally on the screen. Participants were instructed to fixate on the cross or question mark throughout the task. The training phase consisted of 6 blocks of 10 trials, each, with 5 Figure and 5 No-figure stimuli in each block, presented in a pseudorandom order. At the end of each block, summary feedback about accuracy was provided to participants. In the first block, Figure coherence level was 18, decreasing by 2 in each subsequent block (ending at a coherence level of 8). Blocks in the training phase were repeated until the participant’s accuracy was higher than 50% in at least one of the last three blocks (with Figure coherence levels of 12, 10, or 8). Repeated blocks started from the last block with higher than 50% accuracy.

#### Threshold detection setting up the low and the high noise conditions

After training, each participant performed two adaptive threshold detection tasks (~15 mins) to determine the stimuli parameters corresponding to 85% (termed the low-noise (LN) condition) and 65% accuracy (the high-noise (HN) condition) in the figure detection task. In both threshold detection tasks, the trial structure was as described for the training phase, except for the lack of feedback at the end of each trial. As in the training phase, participants were instructed to indicate the presence or absence of a figure in the stimulus, with an emphasis on accuracy.

In the first threshold detection task, the goal was to determine the participant’s individual Figure coherence level that corresponded to ca. 85% accuracy, while keeping the overall number of tones in each cord constant at 20. In the second task, the coherence level was kept constant at the level determined in the LN threshold detection task, and the number of background tones in the chord was increased until performance dropped to ca. 65% accuracy.

In both cases, thresholds were estimated using the QUEST procedure (56), an adaptive staircase method that sets the signal-to-noise ratio (SNR) of the next stimulus to the most probable level of the threshold, as estimated by a Bayesian procedure taking into account all past trials. SNR was determined as the ratio of the number of figure and background tones. Both tasks consisted of one block of 80 trials, with 20 trials added if the standard deviation of the threshold estimate was larger than the median difference between successive SNR levels allowed by FG stimuli parameters. The thresholding phase yielded stimulus parameters corresponding to 65 and 85% accuracy, separately for each participant. Thus, in the main experiment, the LN and HN condition tasks, separately, posed similar levels of difficulty to each participant. The exact parameters used for the QUEST procedure can be found in the GitHub repository of the experiment (https://github.com/dharmatarha/SFG_aging_study/tree/master/threshold).

#### Main figure detection task

In the main part of the experiment (~90 mins), the trial structure and the instructions were identical to those used in the threshold detection procedure. Two conditions (LN and HN) were administered, resulting in four types of stimuli: Figure - LN, No-figure - LN, Figure - HN, and No-figure - HN. Participants received 200 repetitions of each stimulus type for a total of 800 trials. Trials were divided into 10 stimulus blocks of 80 trials each, with each block containing an equal number (20–20) of all four stimulus types in a randomized order. Summary feedback on performance (overall accuracy) was provided to participants after each block. Short breaks were inserted between successive stimulus blocks with additional longer breaks after the 4th and 7th blocks.

### Analysis of behavioral data

From the threshold detection tasks, we analyzed the participants’ coherence level in the LN condition, the number of additional background tones in the HN condition, and log-transformed SNR values for both conditions. A mixed-model ANOVA with the within-subject factor NOISE (LN vs HN) and the between-subject factor GROUP (young adult, normal-hearing elderly, hearing-impaired elderly) was conducted on SNR. For coherence levels (LN only) and the number of additional background tones (HN only), one-way ANOVAs were performed with the between-subject factor GROUP (young, normal-hearing old, hearing-impaired old).

From the main task, detection performance was assessed by the sensitivity index (57), false alarm rate (FA), and mean reaction times (RT). Mixed-mode ANOVAs were conducted with the within-subject factor NOISE, and the between-subject factor GROUP, separately on d’, FA, and RT. Statistical analyses were carried out in Matlab (R2017a). The alpha level was set at 0.05 for all tests. Partial eta squared (*η_p_^2^*) is reported as effect size. Post-hoc pairwise comparisons were computed by Tukey HSD tests.

Pearson’s correlations were calculated between the average of pure-tone audiometry thresholds in the 250 - 8000 Hz range and working memory measures (capacity and control) on one side and behavioral variables from the threshold detection and the figure detection tasks on the other side. Bonferroni correction was used to reduce the potential errors resulting from multiple comparisons.

### Analysis of EEG data

#### EEG recording and preprocessing

EEG was recorded with a Brain Products actiCHamp DC 64-channel EEG system and actiCAP active electrodes. Impedances were kept below 15 kΩ. The sampling rate was 1 kHz with a 100 Hz online low-pass filter applied. Electrodes were placed according to the International 10/20 system with FCz serving as the reference. Eye movements were monitored with bipolar recording from two electrodes placed lateral to the outer canthi of the eyes.

EEG was preprocessed with the EEGlab 11.0.3.1.b toolbox ((58) implemented in Matlab 2018b. Signals were band-pass filtered between 0.5 and 80 Hz using a finite impulse response (FIR) filter (Kaiser windowed, with Kaiser β = 5.65326 and filter length n = 18112). A maximum of two malfunctioning EEG channels were interpolated using the default spline interpolation algorithm implemented in EEGlab. The Infomax algorithm of Independent Component Analysis (ICA) was employed for artifact removal ((58). ICA components from blink artifacts were removed after visually inspecting their topography and the spectral contents of the components. No more than 10 percent of the overall number of ICA components (for a maximum of n = 3) were removed.

#### Event-related brain activity analysis

Epochs were extracted from the continuous EEG records between −800 and +2300 ms relative to the onset of the Figure in Figure trials. For No-figure trials, onsets were selected randomly from the set of Figure onsets in the Figure trials (each Figure onset value from Figure trials was selected only once for a No-figure trial). Only epochs from trials with a correct response (hit for Figure and correct rejection for No-figure trials) were further processed. Baseline correction was applied by averaging voltage values in the [−800 - 0] ms time window. Epochs exceeding the threshold of +/-100 μV change throughout the whole epoch measured at any electrode were rejected. The remaining epochs were averaged separately for each stimulus type and group. The mean number of valid epochs (collapsed across groups) were Figure - HN: 107.78; No-figure - HN: 175.08; Figure - LN: 153.04; No-figure - LN: 173.29. Brain activity within the time windows corresponding to the ORN and P400 ERP components were measured separately for each stimulus type/condition/group. Time windows were defined by visual inspection of grand average ERPs.

The predominantly fronto-central ORN ((25, 30, 31, 48) amplitudes were measured as the average signal in the 250-350 ms latency range from Figure onset for the young adult group and the 350-550 ms latency range for the normal-hearing and hearing-impaired elderly groups at the C3, Cz, and C4 leads. The predominantly parietal P400 ((25, 30, 31, 48) amplitudes were measured as the average signal in the 650-850 ms latency range from the P3, Pz, and P4 electrodes in all three groups.

Peak latency was measured as the latency value of the maximal amplitude within the latency range of ORN and P400 respectively.

The effects of age were tested with mixed mode ANOVAs with within-subject factors FIGURE (Figure vs No-figure), NOISE (LN vs. HN), LATERALITY (left vs midline vs right), and the between-subject factor AGE (young adult vs normal-hearing elderly) on the two ERP amplitudes and peak latencies. Similar mixed-mode ANOVAs were conducted to test the effects of hearing impairment by exchanging AGE for the between-subject factor HEARING IMPAIRMENT (normal-hearing vs hearing-impaired older adults). Post-hoc pairwise comparisons were computed by Tukey HSD tests.

#### EEG source localization

The Brainstorm toolbox (version 2022 January; (59) was used to perform EEG source reconstruction, following the protocol of previous studies (60–62). The MNI/Colin27 brain template was segmented based on the default setting and was entered, along with default electrode locations, into the forward boundary element head model (BEM) provided by the openMEEG algorithm ((63). For the modeling of time-varying source signals (current density) of all cortical voxels, a minimum norm estimate inverse solution was employed using dynamical Statistical Parametric Mapping normalization (64), separately on Figure and No-Figure trials of the LN and HN conditions, and the three groups.

Contrasts were evaluated on the average signal for the time window of interest (250–350 ms for young adults and 350-550 ms for normal-hearing and hearing-impaired older listeners), between Figure trials of the LN and HN conditions, separately for each group by a permutation-based (N=1000) paired sample t-test (alpha level=0.01).

## Acknowledgments

This work was funded by the Hungarian National Research Development and Innovation Office (ANN131305 and PD123790 to BT and K132642 to IW) and the János Bolyai Research Grant awarded to BT (BO/00237/19/2). All data available at https://osf.io/q8n9k/

## Abbreviations

ERP: event-related brain potential
ORN: object-related negativity response
LN: low noise
HN: high noise
SNR: signal-to-noise ratio
IPS: intraparietal sulcus
HT: hearing thresholds
ITI: inter-trial-interval
FA: false alarm rate
RT: mean reaction time
FIR: finite impulse response
IC: independent component analysis
BEM: boundary element head model

