## Supplementary Materials for "The effects of aging and hearing impairment on listening in noise"

**EEG results not involving the GROUP (AGE or HEARING IMPAIRMENT) or NOISE factor**

ORN

*AGE effect on the ORN amplitude.* There was a main effect of FIGURE ($F(1,31)$ = 72.7, $p < 0.001$, $\eta_p^2 = 0.70$) with Figure trials eliciting stronger ORN response (more negative signal) than No-Figure trials and LATERALITY ($F(2,62) = 9.08$, $p < 0.001$, $\eta_p^2 = 0.23$). Post-hoc pairwise comparisons revealed that the ORN was dominant on the left side as the amplitude at C3 amplitudes was larger than at C4 or Cz (p < 0.001, both).

*HEARING IMPAIRMENT effect on the ORN amplitude.* There was a main effect of FIGURE ($F(1,27) = 30.47$, $p < 0.001$, $\eta_p^2 = 0.53$) with Figure trials eliciting stronger ORN response than No-Figure trials and LATERALITY ($F(2,54) = 6.77$, $p < 0.05$, $\eta_p^2 = 0.20$). The significant interaction between FIGURE and LATERALITY ($F(2, 54) = 5.5185$, $p = .007$) was caused by the amplitude at C3 being significantly higher than at Cz for Figure ($p <$ 0.001) but not for No-Figure trials.

*AGE and HEARING IMPAIRMENT effects on the ORN peak latency.*

**P400**

*AGE effects on the P400 amplitude.* There was a main effect of FIGURE ($F(1,31)=77.76$, $p<0.001$, $\eta^2=0.71$) with Figure trials eliciting stronger P400 response (more positive signal) than No-Figure trials and LATERALITY ($F(2,62)=14.71$, $p<0.001$, $\eta_p^2=0.32$). The significant interaction between FIGURE and LATERALITY ($F(2, 62)=4.3340$, $p=.01731$ $\eta_p^2=0.122$) was caused by the amplitude at Pz being significantly higher than at P3 or P4 for Figure ($p < 0.001$, both) but not for No-Figure trials.

*HEARING IMPAIRMENT effects on the P400 amplitude.* There was a main effect of FIGURE ($F(1,27)=55.60$, $p<0.001$, $\eta_p^2=0.67$) with Figure trials eliciting stronger P400 response than No-Figure trials and LATERALITY ($F(2,54)=17.15$, $p<0.001$, $\eta_p^2=0.39$). The significant interaction between FIGURE and LATERALITY ($F(2, 54)=8.6854$, $p<0.001$) was caused by the amplitude at Pz being significantly higher than at P3 or P4 for Figure ($p <$ ????, both) but not for No-Figure trials.

***Supplementary Table 1:*** *Main effects of GROUP on pure tone hearing thresholds in audiometry*

| | *Left Ear* | *Right ear* |
|---|---|---|
| *250 Hz* | *ns.* | *ns.* |
| *500 Hz* | *ns.* | *F(2,46)=3.299, p=0.0458, η2=0.125* |

| | | |
|---|---|---|
| *1000 Hz* | *F(2,46)=19.792, p<0.001, η2=0.462* | *F(2,46)=22.746, p<0.001, η2=0.497* |
| *2000 Hz* | *F(2,46)=19.392, p<0.001, η2=0.458* | *F(2,46)=27.845, p<0.001, η2=0.548* |
| *4000 Hz* | *F(2,46)=28.353, p<0.001, η2=0.552* | *F(2,46)=34.555, p<0.001, η2=0.601* |
| *6000 Hz* | *F(2,46)=69.681, p<0.001, η2=0.752* | *F(2,46)=60.194, p<0.001, η2=0.723* |
| *8000 Hz* | *F(2,46)=121.157, p<0.001, η2=0.841* | *F(2,46)=133.492, p<0.001, η2=0.853* |

***Supplementary Table 2:*** *The subset of items from the HHIE administered in the study*

| |
|---|
| *1. Does a hearing problem cause you to feel embarrassed when you meet new people?* |
| *2. Does a hearing problem cause you to feel frustrated when talking to members of your family?* |
| *3. Do you have difficulty hearing when someone speaks in a whisper?* |
| *4. Do you feel handicapped by a hearing problem?* |
| *5. Does a hearing problem cause you difficulty when visiting friends, relatives, or neighbors?* |
| *6. Does a hearing problem cause you to attend religious services less often than you would like?* |
| *7. Does a hearing problem cause you to have arguments with family members?* |
| *8. Does a hearing problem cause you to have difficulty when listening to television or radio?* |
| *9. Do you feel that any difficulty with your hearing limits/hampers your personal or social life?* |
| *10. Does a hearing problem cause you difficulties when in restaurant?* |
